# Differential Expression Analysis for RNAseq using Poisson Mixed Models

**DOI:** 10.1101/073403

**Authors:** Shiquan Sun, Michelle Hood, Laura Scott, Qinke Peng, Sayan Mukherjee, Jenny Tung, Xiang Zhou

## Abstract

Identifying differentially expressed (DE) genes from RNA sequencing (RNAseq) studies is among the most common analyses in genomics. However, RNAseq DE analysis presents several statistical and computational challenges, including over-dispersed read counts and, in some settings, sample non-independence. Previous count-based methods rely on simple hierarchical Poisson models (e.g., negative binomial) to model independent over-dispersion, but do not account for sample non-independence due to relatedness, population structure and/or hidden confounders. Here, we present a Poisson mixed model with two random effects terms that account for both independent over-dispersion and sample non-independence. We also develop a scalable sampling-based inference algorithm using a latent variable representation of the Poisson distribution. With simulations, we show that our method properly controls for type I error and is generally more powerful than other widely used approaches, except in small samples (n<15) with other unfavorable properties (e.g., small effect sizes). We also apply our method to three real data sets that contain related individuals, population stratification, or hidden confounders. Our results show that our method increases power in all three data compared to other approaches, though the power gain is smallest in the smallest sample (n=6). Our method is implemented in MACAU, freely available at www.xzlab.org/software.html.

## Introduction

RNA sequencing (RNAseq) has emerged as a powerful tool for transcriptome analysis, thanks to its many advantages over previous microarray techniques (1-3). Compared with microarrays, RNAseq has increased dynamic range, does not rely on *a priori-chosen* probes, and can thus identify previously unknown transcripts and isoforms. It also yields allelic-specific expression estimates and genotype information inside expressed transcripts as a useful by-product (4-7). Because of these desirable features, RNAseq has been widely applied in many areas of genomics and is currently the gold standard method for genome-wide gene expression profiling.

One of the most common analyses of RNAseq data involves identification of differentially expressed (DE) genes. Identifying DE genes that are influenced by predictors of interest -- such as disease status, risk factors, environmental covariates, or genotype -- is an important first step towards understanding the molecular basis of disease susceptibility as well as the genetic and environmental basis of gene expression variation. Progress towards this goal requires statistical methods that can handle the complexities of the increasingly large and structurally complex RNAseq data sets that are now being collected from population and family studies (8,9). Indeed, even in classical treatment-control comparisons, the importance of larger sample sizes for maximizing power and reproducibility is increasingly well appreciated (10,11). However, identifying DE genes from such studies presents several key statistical and computational challenges, including accounting for ambiguously mapped reads (12), modeling uneven distribution of reads inside a transcript (13), and inferring transcript isoforms (14).

A fundamental challenge shared by all DE analyses in RNAseq, though, is accounting for the count nature of the data (3,15,16). In most RNAseq studies, the number of reads mapped to a given gene or isoform (following appropriate data processing and normalization) is often used as a simple and intuitive estimate of its expression level (13,14,17). As a result, RNAseq data display an appreciable dependence between the mean and variance of estimated gene expression levels: highly expressed genes tend to have high read counts and subsequently high between-sample variance, and vice versa (15,18). To account for the count nature of the data and the resulting mean-variance dependence, most statistical methods for DE analysis model RNAseq data using discrete distributions. For example, early studies showed that gene expression variation across technical replicates can be accurately described by a Poisson distribution (19-21). More recent methods also take into account over-dispersion across biological replicates (22,23) by replacing Poisson models with negative binomial models (15,16,24-28) or other related approaches (18, 29-32). While non-count based methods are also commonly used (primarily relying on transformation of the count data to more flexible, continuous distributions (33,34)), recent comparisons have highlighted the benefits of modeling RNAseq data using the original counts and accounting for the resulting mean-variance dependence (35-38), consistent with observations from many count data analyses in other statistical settings (39). Indeed, accurate modeling of mean-variance dependence is one of the keys to enable powerful DE analysis with RNAseq, especially in the presence of large sequencing depth variation across samples (25,33,40).

A second important feature of many RNAseq data sets, which has been largely overlooked in DE analysis thus far, is that samples often are not independent. Sample non-independence can result from individual relatedness, population stratification, or hidden confounding factors. For example, it is well known that gene expression levels are heritable. In humans, the narrow-sense heritability of gene expression levels averages from 15%-34% in peripheral blood (41-45) and is about 23% in adipose tissue (41), with a maximum heritability in both tissues as high as 90% (41,42). Similarly, in baboons, gene expression levels are about 28% heritable in the peripheral blood (7). Some of these effects are attributable to nearby, putatively *cis*-acting genetic variants: indeed, recent studies have shown that the expression levels of almost all genes are influenced by cis-eQTLs and/or display allelic specific expression (ASE) (3,7,46-48). However, the majority of heritability is often explained by distal genetic variants (i.e., *trans-*QTLs, which account for 63%-84% of heritability in humans (41) and baboons (7)). Because gene expression levels are heritable, they will covary with kinship or population structure. Besides kinship or population structure, hidden confounding factors, commonly encountered in sequencing studies (49-52), can also induce similarity in gene expression levels across many genes even when individuals are unrelated (53-57). Failure to account for this gene expression covariance due to sample non-independence could lead to spurious associations or reduced power to detect true DE effects. This phenomenon has been extensively documented in genome-wide association studies (9,58,59) and more recently, in bisulfite sequencing studies (60), but is less explored in RNAseq studies. In particular, none of the currently available count-based methods for identifying DE genes in RNAseq can appropriately control for sample non-independence. Consequently, even though count-based methods have been shown to be more powerful, recent RNAseq studies have turned to linear mixed models, which are specifically designed for quantitative traits, to deal with the confounding effects of kinship, population structure, or hidden confounders (7,42,61).

Here, we present a Poisson mixed model (PMM) that can explicitly model both over-dispersed count data and sample non-independence in RNAseq data for effective DE analysis. To make our model scalable to large data sets, we also develop an accompanying efficient inference algorithm based on an auxiliary variable representation of the Poisson model (62-64) and recent advances in mixed model methods (9,59,65). We refer to the combination of the statistical method and the computational algorithm developed here as MACAU (Mixed model Association for Count data via data AUgmentation), which effectively extends our previous method of the same name on the simpler binomial model (60) to the more difficult Poisson model. MACAU works directly on RNAseq count data and introduces two random effects terms to both control for sample non-independence and account for additional independent over-dispersion. As a result, MACAU properly controls for type I error in the presence of sample non-independence and, in a variety of settings, is more powerful for identifying DE genes than other commonly used methods. We illustrate the benefits of MACAU with extensive simulations and real data applications to three RNAseq studies.

## Methods and Materials

### Methods for Comparison

We compared the performance of seven different methods in the main text: (1) our Poisson mixed model implemented in the MACAU software package (60); (2) the linear model implemented in the *lm* function in R; (3) the linear mixed model implemented in the GEMMA software package (9,59,66); (4) the Poisson model implemented in the *glm* function in R; (5) the negative binomial model implemented in the *glm.nb* function in R; (6) edgeR implemented in the *edgeR* package in R (25); (7) DESeq2 implemented in the *DESeq2* package in R (24). All methods were used with default settings. The performance of each method in simulations was evaluated using the area under the curve (AUC) function implemented in the *pROC* package in R (67), a widely used benchmark for RNAseq method comparisons (68).

Both the linear model and the linear mixed model require quantitative phenotypes. Here, we considered six different transformations of count data to quantitative values, taking advantage of several methods proposed to normalize RNAseq data (e.g., (12-14,17,22,33,69)): (1) quantile normalization (TRCQ), where we first divided the number of reads mapped to a given gene by the total number of read counts for each individual, and then for each gene, quantile normalized the resulting proportions across individuals to a standard normal distribution (7); (2) total read count normalization (TRC), where we divided the number of reads mapped to a given gene by the total number of read counts for each individual (i.e. CPM, counts per million; without further transformation to a standard normal within genes: (25)); (3) upper quantile normalization (UQ), where we divided the number of reads mapped to a given gene by the upper quantile (75-th percentile) of all genes for each individual (70); (4) relative log expression normalization (RLE) (15); (5) the trimmed mean of M-values (TMM) method (40) where we divided the number of reads mapped to a given gene by the normalization factor output from TMM; and (6) VOOM normalization (33). Simulations in a supplementary figure showed that TRCQ, VOOM and TRC worked better than the other three methods, with TRCQ showing a small advantage. Therefore, we report results using TRCQ throughout the text.

### Simulations

To make our simulations as realistic as possible, we simulated the gene expression count data based on parameters inferred from a real baboon data set that contains 63 samples (see the next section for a detailed description of the data). We varied the sample size (*n*) in the simulations (*n* = 6, 10, 14, 63, 100, 200, 500, 800, or 1000). For *n* = 63, we used the baboon relatedness matrix ***K*** (7). For sample simulations with *n* > 63, we constructed a new relatedness matrix ***K*** by filling in its off-diagonal elements with randomly drawn off-diagonal elements from the baboon relatedness matrix following (60). For sample simulations with *n* < 63, we constructed a new relatedness matrix ***K*** by randomly sub-sampling individuals from the baboon relatedness matrix. In cases where the resulting ***K*** was not positive definite, we used the *nearPD* function in R to find the closest positive definite matrix as the final ***K***. In most cases, we simulated the total read count *N_i_* for each individual from a discrete uniform distribution with a minimum (=1,770,083) and a maximum (=9,675,989) total read count (i.e. summation of read counts across all genes) equal to the minimum and maximum total read counts from the baboon data. We scaled the total read counts to ensure that the coefficient of variation was small (CV = 0.3), moderate (CV = 0.6) or high (CV = 0.9) across individuals (i.e. *N_new_* = *N̅ +* (*N − N̅*) *CV sd*(*N*) */N*), and then discretized them. In the special case where CV = 0.3 and *n* = 63, we directly used the observed total read counts per individual *i* (*N_i_*) from the baboon data (which has a CV = 0.33).

We then repeatedly simulated a continuous predictor variable *x* from a standard normal distribution (without regard to the pedigree structure). We estimated the heritability of the continuous predictor using GEMMA, and retained *x* if the heritability (*h_x_^2^*) estimate (with ± 0.01 tolerance) was 0, 0.4 or 0.8, representing no, moderate and highly heritable predictors. Using this procedure, approximately 30 percent of *x* values generated were retained, with different retention percentages for different heritability values.

Based on the simulated sample size, total read counts and continuous predictor variable, we simulated gene expression values using the following procedure. For the expression of each gene in turn, we simulated the genetic random effects ***g*** from a multivariate normal distribution with covariance ***K***. We simulated the environmental random effects ***e*** based on independent normal distributions. We scaled the two sets of random effects to ensure a fixed value of heritability (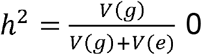 or 0.3 or 0.6) and a fixed value of over-dispersion variance (*σ^2^* = *V*(*g*) *+ V*(*e*) = 0.1, 0.25 or 0.4, close to the lower, median and upper quantiles of the over-dispersion variance inferred from the baboon data, respectively), where the function V(∙) denotes the sample variance. We then generated the effect size *β* of the predictor variable on gene expression. The effect size was either 0 (for non-DE genes) or generated to explain a certain percentage of variance in log(*λ*) (i.e. 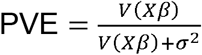; for DE genes). PVE values were 15%, 20%, 25%, 30% or 35% to represent different effect sizes. The predictor effects *Xβ,* genetic effects ***g***, environmental effects ***e***, and an intercept (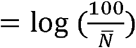) to ensure that the expected simulated count is 100) were then summed together to yield the latent variable log(*λ*) = *μ*+*Xβ*+*g + e.* Note that *h^2^* does not include the contribution of *Xβ,* which in many cases represent non-genetic effects. Finally, the read counts were simulated based on a Poisson distribution with rate determined by the total read counts and the latent variable λ, or *y_i_~Poi*(*N_i_λ_i_*) for the *i*'th individual.

With the above procedure, we first simulated data for *n* = 63, CV = 0.3, *h_x_^2^* = 0, PVE = 0.25, *h^2^* = 0.3 and *σ^2^* = 0.25. We then varied one parameter at a time to generate different scenarios for comparison. In each scenario, conditional on the sample size, total read counts, and continuous predictor variable, we performed 10 simulation replicates, where “replication” is at the level described in the paragraph above. Each replicate consisted of 10,000 genes. For examining type I error control, all 10,000 genes were non-DE. For the power comparison, 1,000 genes were DE while 9,000 were non-DE.

### RNAseq Data Sets

We considered three published RNAseq data sets in this study, which include small (n<15), medium (15≤n≤100), and large (n>100) sample sizes (based on current RNAseq sample sizes in the literature).

The first RNAseq data set was collected from blood samples of yellow baboons (7) from the Amboseli ecosystem of southern Kenya as part of the Amboseli Baboon Research Project (ABRP) (71). The data are publicly available on GEO with accession number GSE63788. Read counts were measured on 63 baboons and 12,018 genes after stringent quality control as in (7). As in (7), we computed pairwise relatedness values from previously collected microsatellite data (72,73) using the software COANCESTRY (74). The data contains related individuals: 16 pairs of individuals have a kinship coefficient exceeding 1/8 and 48 pairs exceed 1/16. We obtained sex information for each individual from GEO. Sex differences in health and survival are major topics of interest in medicine, epidemiology, and evolutionary biology (72,75). Therefore, we used this data set to identify sex-related gene expression variation. In the analysis, we included the top 5 expression PCs as covariates to control for potential batch effects following the original study (7).

The second RNAseq data set was collected from skeletal muscle samples of Finnish individuals (61) as part of the FUSION project (76,77). The data are publicly available in dbGaP with accession code phs001068.v1.p1. Among the 271 individuals in the original study, we selected 267 individuals who have both genotypes and gene expression measurements. Read counts were obtained on these 267 individuals and 21,753 genes following the same stringent quality control as in the FUSION study. For genotypes, we excluded SNPs with minor allele frequency (MAF) < 0.05 and Hardy-Weinberg equilibrium *p*-value < 10^−6^. We used the remaining 5,696,681 SNPs to compute the relatedness matrix using GEMMA. The data contains remotely related individuals (3 pairs of individuals have a kinship coefficient exceeding 1/32 and 6 pairs exceed 1/64) and is stratified by the municipality from which samples were collected. Two predictors from the data were available to us: the oral glucose tolerance test (OGTT) which classifies *n* = 162 individuals as either T2D patient (n = 66) or normal glucose tolerance (NGT; i.e., control, *n* = 96); and a T2D-related quantitative trait -- fasting glucose levels (GL) -- measured on all *n* = 267 individuals. We used these data to identify genes whose expression level is associated with either T2D or GL. In the analysis, we included age, sex and batch labels as covariates following the original study (61).

The third RNAseq data set was collected from lymphoblastoid cell lines (LCLs) derived from 69 unrelated Nigerian individuals (YRI) (3). The data are publicly available on GEO with accession number GSE19480. Following the original study (3), we aligned reads to the human reference genome (version hg19) using BWA (78). We counted the number of reads mapped to each gene on either autosomes or the X chromosome using Ensembl gene annotation information obtained from the UCSC genome browser. We then filtered out lowly expressed genes with zero counts in over 90% of individuals. In total, we obtained gene expression measurements on 13,319 genes. Sex is the only phenotype available in the data and we used sex as the predictor variable to identify sex-associated genes. To demonstrate the efficacy of MACAU in small samples, we randomly subsampled individuals from the data to create small data sets with either *n* = 6 (3 males and 3 females) or *n* = 10 (5 males and 5 females), or *n* = 14 individuals (7 males and 7 females). For each sample size n, we performed 20 replicates of subsampling and we evaluated method performance by averaging across these replicates. In each replicate, following previous studies (53-57), we used the gene expression covariance matrix as ***K*** (i.e. ***K*** *=* ***XX^T^****/***p**, where ***X*** is the normalized gene expression matrix and *p* is the number of genes) and applied MACAU to identify sex-associated genes. Note that the gene expression covariance matrix ***K*** contains information on sample non-independence caused by hidden confounding factors (53-57), and by incorporating ***K***, MACAU can be used to control for hidden confounding factors that are commonly observed in sequencing data sets (49-52).

For each of these RNAseq data sets and each trait, we used a constrained permutation procedure to estimate the empirical false discovery rate (FDR) of a given analytical method. In the constrained permutation procedure, we permuted the predictor across individuals, estimated the heritability of the permuted predictor, and retained the permutation only if the permuted predictor had a heritability estimate (*h_x_^2^*) similar to the original predictor with ± 0.01 tolerance (for the original predictors: *h_x_^2^* = 0.0002 for sex in the baboon data; *h_x_^2^* = 0.0121 for T2D and *h_x_^2^* = 0.4023 for GL in the FUSION data; *h_x_^2^* are all close to zero with small variations depending on the sub-sample size in the YRI data). We then analyzed all genes using the permuted predictor. We repeated the constrained permutation procedure and analysis 10 times, and combined the *p*-values from these 10 constrained permutations. We used this set of *p*-values as a null distribution from which to estimate the empirical false discovery rate (FDR) for any given *p*-value threshold (60). This constrained procedure thus differs from the usual unconstrained permutation procedure (every permutation retained) (79) in that it constrains the permuted predictor to have the same *h_x_^2^* as the original predictor. We chose to use the constrained permutation procedure here because the unconstrained procedure is invalid under the mixed model assumption: the subjects are not exchangeable in the presence of sample non-independence (individual relatedness, population structure, or hidden confounders) (79,80). To validate our constrained permutation procedure and test its effectiveness in estimating FDR, we performed a simulation with 1,000 DE genes and 9,000 non-DE genes as described above. We considered three predictor variables *x* with different heritability: *h_x_^2^* = 0, *h_x_^2^* = 0.4, and *h_x_^2^* = 0.8. For each predictor variable and each *p*-value threshold, we computed the true FDR and then estimated the FDR based on either the constrained or unconstrained permutation procedures. The simulation results presented in a supplementary figure demonstrate that the constrained permutation procedure provides a much more accurate estimate of the true FDR while the unconstrained permutation procedure often under-estimates the true FDR. Therefore, we applied the constrained permutation procedure for all real data analysis.

Finally, we investigated whether the methods we compared were sensitive to outliers (31,81,82) in the first two data sets. To examine outlier sensitivity, we first identified genes with potential outliers using BBSeq (18). In total, we identified 8 genes with potential outliers in the baboon data, 130 genes with potential outliers in the FUSION data (*n* = 267) and 43 genes with potential outliers in the subset of the FUSION data for which we had T2D diagnoses (*n* = 162). We counted the number of genes with potential outliers in the top 1,000 genes with strong DE association evidence. In the baboon data, 4 genes with potential outliers are in the top 1,000 genes with the strongest sex association determined by various methods: 2 of them by the negative binomial model, 3 of them by the Poisson model, but 0 of them by MACAU, linear model, or GEMMA. In the FUSION data, for T2D analysis, 9 genes with potential outliers are in the top 1,000 genes with the strongest T2D association determined by various methods: 1 by MACAU, 3 by negative binomial, 6 by Poisson, 1 by linear, and 1 by GEMMA. For GL analysis, 15 genes with potential outliers are in the top 1,000 genes with the strongest GL association determined by various methods: 2 by MACAU, 7 by negative binomial, 9 by Poisson, 3 by linear, and 3 by GEMMA. All outliers are presented in supplementary figures. Therefore, the influence of outliers on DE analysis is small in the real data.

## Results

### MACAU Overview

Here, we provide a brief overview of the Poisson mixed model (PMM); more details are available in the Supplementary Material. To identify DE genes with RNAseq data, we examine one gene at a time. For each gene, we model the read counts with a Poisson distribution

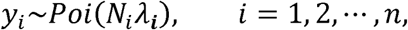

where for the *i*'th individual, *y_i_* is the number of reads mapped to the gene (or isoform); *N_i_* is the total read counts for that individual summing read counts across all genes; and *λ_i_* is an unknown Poisson rate parameter. We model the log-transformed rate *λ_i_* as a linear combination of several parameters

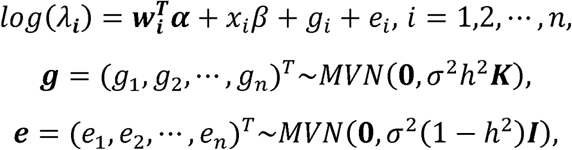

where ***w***_*i*_ is a *c*-vector of covariates (including the intercept); ***α*** is a c-vector of corresponding coefficients; *x_i_* represents the predictor variable of interest (e.g. experimental perturbation, sex, disease status, or genotype); *β* is its coefficient; ***g*** is an *n*-vector of genetic effects; ***e*** is an *n*-vector of environmental effects; ***K*** is an *n* by *n* positive semi-definite matrix that models the covariance among individuals due to individual relatedness, population structure, or hidden confounders; ***I*** is an *n* by *n* identity matrix that models independent environmental variation; *σ^2^h^2^* is the genetic variance component; *σ*^2^(*1−h*^2^) is the environmental variance component; and *MVN* denotes the multivariate normal distribution. In the above model, we assume that ***K*** is known and can be computed based on either pedigree, genotype, or the gene expression matrix (9). For pedigree/genotype data, when ***K*** is standardized to have ***tr***(***K***)*/n =* 1*, h^2^* ∈ [0,1] has the usual interpretation of heritability (9), where the tr(∙) denotes the trace of a matrix. Importantly, unlike several other DE methods (15,25), our model can deal with both continuous and discrete predictor variables.

Both of the random effects terms ***g*** and ***e*** model over-dispersion, the extra variance not explained by a Poisson model. However, the two terms ***g*** and ***e*** model two different aspects of over-dispersion. Specifically, ***g*** models the fraction of the extra variance that is explained by sample non-independence while ***e*** models the fraction of the extra variance that is independent across samples. For example, let us consider a simple case in which all samples have the same sequencing depth (i.e. *N_i_* = *N)* and there is only one intercept term *μ* included as the covariate. In this case, the random effects term ***e*** models the independent over-dispersion: without ***g***, *V*(*y*) = *E*(*y*) (1 + *E*(*y*(*e_σ_2__* − 1)) is still larger than the mean *E*(*y* = *Ne_σ+μ_2_/2_*, with the difference between the two increasing with increasing *σ*^2^. In a similar fashion, the random effects term ***g*** models the non-independent over-dispersion by accounting for the sample covariance matrix ***K***. By modeling both aspects of over-dispersion, our PMM effectively generalizes the commonly used negative binomial model -- which only models independent extra variance -- to account for sample non-independence. In addition, our PMM naturally extends the commonly used linear mixed model (LMM) (9,65,83) to modeling count data.

Our goal here is to test the null hypothesis that gene expression levels are not associated with the predictor variable of interest, or *H*_0_:*β* = 0. Testing this hypothesis requires estimating parameters in the PMM (as has previously been done in other settings (84,85), including for modeling uneven RNAseq read distribution inside transcripts (13); details in Supplementary Material). The PMM belongs to the generalized linear mixed model family, where parameter estimation is notoriously difficult because of the random effects and the resulting intractable *n*-dimensional integral in the likelihood. Standard estimation methods rely on numerical integration (86) or Laplace approximation (87,88), but neither strategy scales well with the increasing dimension of the integral, which in our case equals the sample size. As a consequence, standard approaches often produce biased estimates and overly narrow (i.e., anti-conservative) confidence intervals (89-95). To overcome the high-dimensionality of the integral, we instead develop a novel Markov Chain Monte Carlo (MCMC) algorithm, which, with enough iterations, can achieve high inference accuracy (96,97). We use MCMC to draw posterior samples but rely on the asymptotic normality of both the likelihood and the posterior distributions (98) to obtain the approximate maximum likelihood estimate *β̂_j_* and its standard error 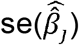. With *β̂_j_* and 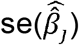, we can construct approximate Wald test statistics and *p*-values for hypothesis testing (Supplementary Material). Although we use MCMC, our procedure is frequentist in nature.

At the technical level, our MCMC algorithm is also novel, taking advantage of an auxiliary variable representation of the Poisson likelihood (62-64) and recent linear algebra innovations for fitting linear mixed models (9,59,65). Our MCMC algorithm introduces *two* continuous latent variables for each individual to replace the count observation, effectively extending our previous approach of using *one* latent variable for the simpler binomial distribution (60). Compared with a standard MCMC, our new MCMC algorithm reduces the computational complexity of each MCMC iteration from cubic to quadratic with respect to the sample size. Therefore, our method is orders of magnitude faster than the popular Bayesian software MCMCglmm (99) and can be used to analyze hundreds of samples and tens of thousands of genes with a single desktop PC (Figure S1). Although our procedure is stochastic in nature, we find the MCMC errors are often small enough to ensure stable *p*-values across independent MCMC runs (Figure S2).

### Simulations: control for sample non-independence

We performed a series of simulations to compare the performance of the PMM implemented in MACAU with four other commonly used methods: (1) a linear model; (2) the linear mixed model implemented in GEMMA (9,59); (3) a Poisson model; and (4) a negative binomial model. We used quantile-transformed data for linear model and GEMMA (see Methods and Materials for normalization details and a comparison between various transformations; Figure S3) and used raw count data for the other three methods. To make our simulations realistic, we use parameters inferred from a published RNAseq data set on a population of wild baboons (7,71) to perform simulations (Methods and Materials); this baboon data set contains known related individuals and hence invokes the problem of sample non-independence outlined above.

Our first set of simulations was performed to evaluate the effectiveness of MACAU and the other four methods in controlling for sample non-independence. To do so, we simulated expression levels for 10,000 genes in 63 individuals (the sample size from the baboon data set). Simulated gene expression levels are influenced by both independent environmental effects and correlated genetic effects, where genetic effects are simulated based on the baboon kinship matrix (estimated from microsatellite data (7)) with either zero (*h*^2^ =0.0), moderate (*h*^2^ = 0.3), or high (*h*^2^ = 0.6) heritability values. We also simulated a continuous predictor variable x that is itself moderately heritable (*h_x_^2^* = 0.4). Because we were interested in the behavior of the null in this set of simulations, gene expression levels were not affected by the predictor variable (i.e., no genes were truly DE).

Figures 1, S4, and S5 show quantile-quantile plots for analyses using MACAU and the other four methods against the null (uniform) expectation, for *h*^2^ = 0.6, *h*^2^ = 0.3, and *h*^2^ = 0.0 respectively. When genes are heritable and the predictor variable is also correlated with individual relatedness, then the resulting *p*-values from the DE analysis are expected to be uniform only for a method that properly controls for sample non-independence. If a method fails to control for sample non-independence, then the *p*-values would be inflated, resulting in false positives.

**Figure 1.**
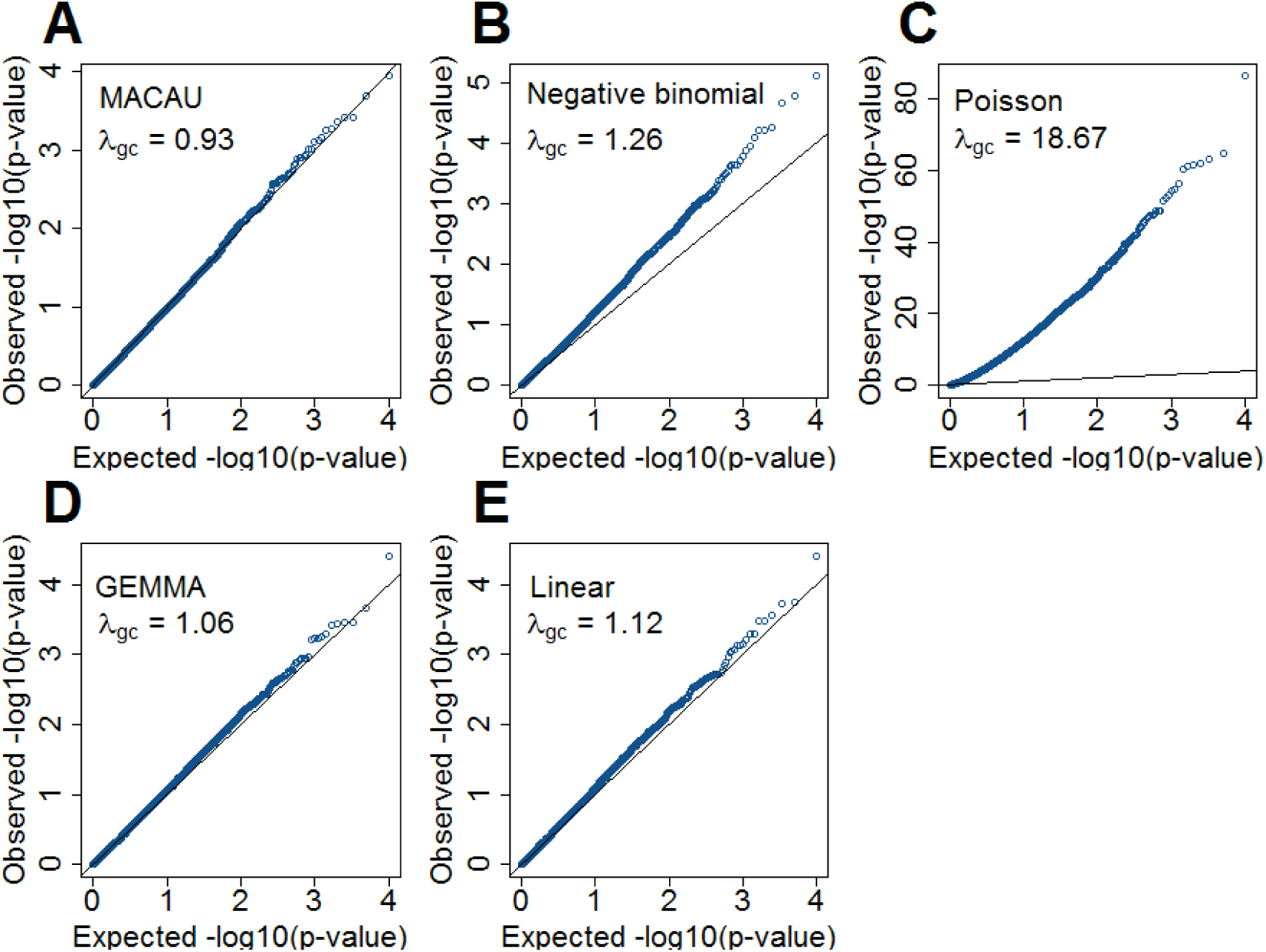
QQ-plots comparing expected and observed *p*-value distributions generated by different methods for the null simulations in the presence of sample non-independence. In each case, 10,000 non-DE genes were simulated with *n* = 63, CV = 0.3, *σ^2^* = 0.25, *h*^2^= 0.6 and *h*_x_^2^ = 0.4. Methods for comparison include MACAU (A), Negative binomial (B), Poisson (C), GEMMA (D), and Linear (E). Both MACAU and GEMMA properly control for type I error well in the presence of sample non-independence. *λ_gc_* is the genomic control factor.

Our results show that, because MACAU controls for sample non-independence, the *p*-values from MACAU follow the expected uniform distribution closely (and are slightly conservative) regardless of whether gene expression is moderately or highly heritable. The genomic control factors from MACAU are close to 1 (Figures 1 and S4). Even if we use a relatively relaxed *q*-value cutoff of 0.2 to identify DE genes, we do not incorrectly identify any genes as DE with MACAU. In contrast, the *p*-values from negative binomial are inflated and skewed towards low (significant) values, especially for gene expression levels with high heritability. With negative binomial, 27 DE genes (when *h*^2^ = 0.3) or 21 DE genes (when *h*^2^ = 0.6) are erroneously detected at the *q*-value cutoff of 0.2. The inflation of *p*-values is even more acute in Poisson, presumably because the Poisson model accounts for neither individual relatedness nor over-dispersion. For non-count-based models, the *p*-values from a linear model are slightly skewed towards significant values, with 3 DE genes (when *h*^2^ = 0.3) and 1 DE gene (when *h*^2^ = 0.6) erroneously detected at *q* < 0.2. In contrast, because the LMM in GEMMA also accounts for individual relatedness, it controls for sample non-independence well. Finally, when genes are not heritable, all methods except Poisson correctly control type I error (Figure S5).

Two important factors influence the severity of sample non-independence in RNAseq data (Figure 2). First, the inflation of *p*-values in the negative binomial, Poisson and linear models becomes more acute with increasing sample size. In particular, when *h_x_^2^* = 0.4, with a sample size of *n =* 1,000, *λ_gc_* from the negative binomial, Poisson and linear models reaches 1.71, 82.28, and 1.41, respectively. In contrast, even when *n* = 1,000, *λ_gc_* from both MACAU and GEMMA remain close to 1 (0.97 and 1.01, respectively). Second, the inflation of *p*-values in the three models also becomes more acute when the predictor variable is more correlated with population structure. Thus, for a highly heritable predictor variable (*h_x_^2^* = 0.8), *λ_gc_* (when *n* = 1,000) from the negative binomial, Poisson and linear models increases to 2.13, 101.43, and 1.81, respectively, whereas *λ_gc_* from MACAU and GEMMA remains close to 1 (1.02 and 1.05).

**Figure 2.**
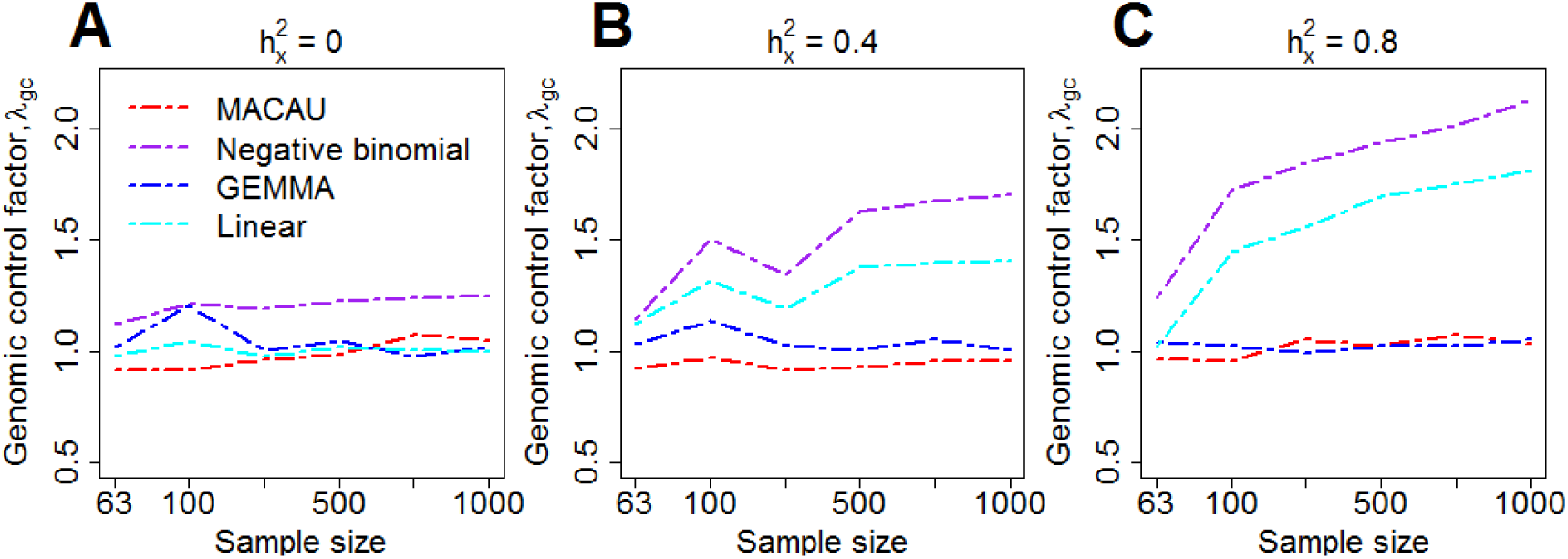
Comparison of the genomic control factor *A_gc_* from different methods for the null simulations in the presence of sample non-independence. 10,000 null genes were simulated with CV = 0.3, *σ*^2^ *=* 0.25, *h*^2^ *=* 0.6, and (A) *h_x_*^2^ = 0; (B) *h_x_*^2^ = 0.4; or (C) *h_x_*^2^ = 0.8. *λ_gc_* (*y*-axis) changes with sample size *n* (*x*-axis). Methods for comparison were MACAU (red), Negative binomial (purple), GEMMA (blue), and Linear (cyan). Both MACAU and GEMMA provide calibrated test statistics in the presence of sample non-independence across a range of settings. *λ_gc_* from Poisson exceeds 10 in all settings and is thus not shown

We also compared MACAU with edgeR (25) and DESeq2 (15), two commonly used methods for DE analysis (38,100). Because edgeR and DESeq2 were designed for discrete predictor valuables, we discretized the continuous predictor *x* into 0/1 based on the median predictor value across individuals. We then applied all methods to the same binarized predictor values for comparison. Results are shown in Figure S6. For the five methods compared above, the results on binarized values are comparable with those for continuous variables (i.e. Figure S6 vs Figure 1). Both edgeR and DESeq2 produce anticonservative *p*-values and perform similarly to the negative binomial model in terms of type I error control.

Finally, we explored the use of principal components (PCs) from the gene expression matrix or the genotype matrix to control for sample non-independence. Genotype PCs have been used as covariates to control for population stratification in association studies (101). However, recent comparative studies have shown that using PCs is less effective than using linear mixed models (83, 102). Consistent with the poorer performance of PCs in association studies (83, 102), using the top PCs from either the gene expression matrix or the genotype matrix does not improve type I error control for negative binomial, Poisson, linear, edgeR or DESeq2 approaches (Figures S7 and S8).

### Simulations: power to identify DE genes

Our second set of simulations was designed to compare the power of different methods for identifying DE genes, again based on parameters inferred from real data. This time, we simulated a total of 10,000 genes, among which 1,000 genes were truly DE and 9,000 were non-DE. For the DE genes, simulated effect sizes corresponded to a fixed proportion of variance explained (PVE) in gene expression levels that ranged from 15% to 35%. For each set of parameters, we performed 10 replicate simulations and measured model performance based on the area under the curve (AUC) (as in (35,68,103)). We also examined several key factors that could influence the relative performance of the alternative methods: (1) gene expression heritability *(h*^2^); (2) correlation between the predictor variable *x* and genetic relatedness (measured by the heritability of *x*, or *h_x_^2^*); (3) variation of the total read counts across samples (measured by the coefficient of variation, or CV); (4) the over-dispersion parameter (*σ^2^*); (5) the effect size (PVE); and (6) sample size (n). To do so, we first performed simulations using a default set of values (*h*^2^ = 0.3, *h_x_^2^* = 0, CV = 0.3, *σ^2^* = 0.25, PVE = 0.25, and *n* = 63) and then varied them one at a time to examine the influence of each factor on the relative performance of each method.

Our results show that MACAU works either as well as or better than other methods in almost all settings (Figures 3, S9-S14), probably because it both models count data directly and controls for sample non-independence. In contrast, the Poisson approach consistently fared the worst across all simulation scenarios, presumably because it fails to account for any sources of over-dispersion (Figures 3, S9-S14).

**Figure 3.**
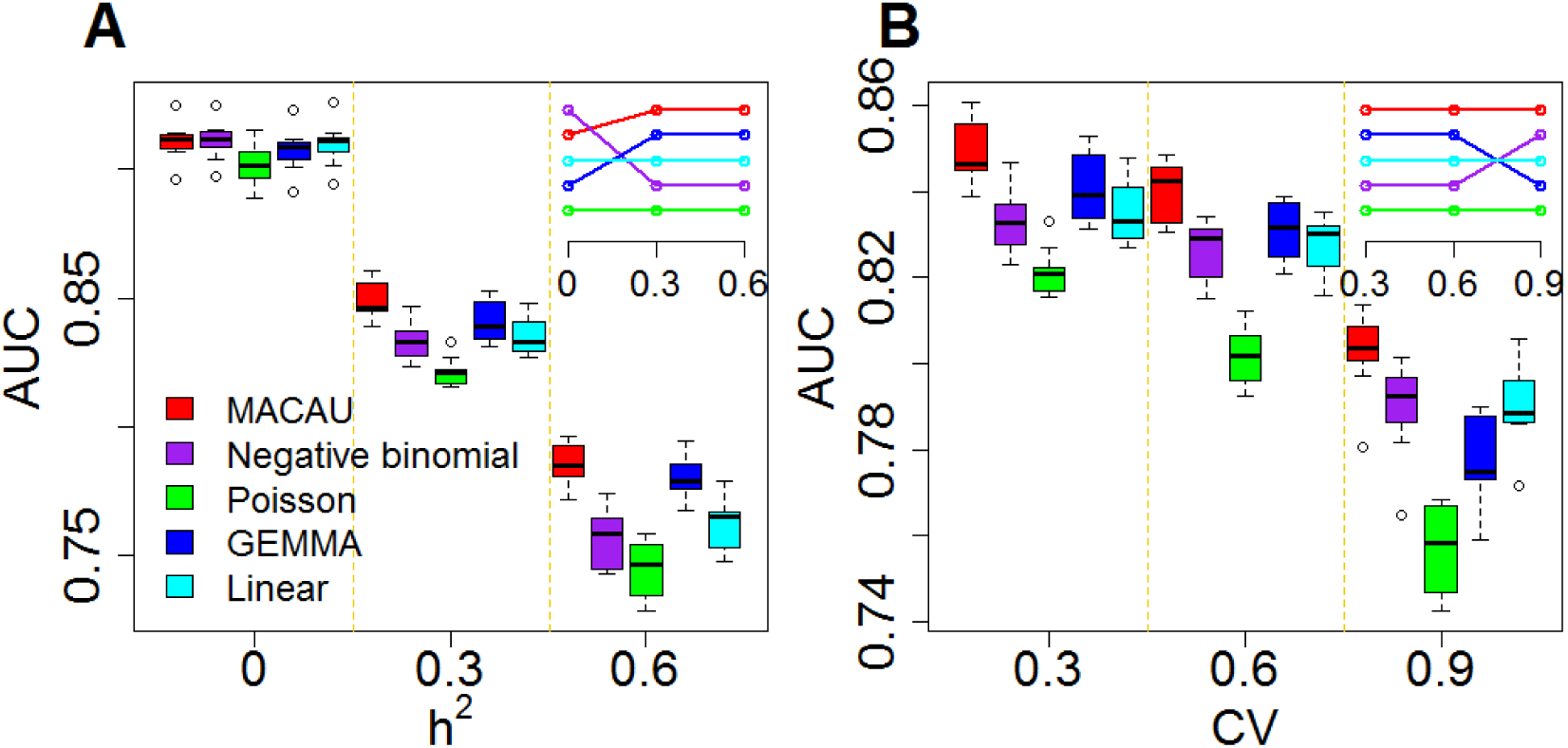
MACAU exhibits increased power to detect true positive DE genes across a range of simulation settings. Area under the curve (AUC) is shown as a measure of performance for MACAU (red), Negative binomial (purple), Poisson (green), GEMMA (blue), and Linear (cyan). Each simulation setting consists of 10 simulation replicates, and each replicate includes 10,000 simulated genes, with 1,000 DE and 9,000 non-DE. We used *n* = 63, *h*_x_^2^ = 0.0, PVE = 0.25, *σ*^2^ = 0.25. In (A) we increased *h*^2^ while maintaining CV = 0.3 and in (B) we increased CV while maintaining *h*^2^ = 0.3. Boxplots of AUC across replicates for different methods show that (A) heritability (*h*^2^) influences the relative performance of the methods that account for sample non-independence (MACAU and GEMMA) compared to the methods that do not (negative binomial, Poisson, linear); (B) variation in total read counts across individuals, measured by the coefficient of variation (CV), influences the relative performance of GEMMA and negative binomial. Insets in the two figures show the rank of different methods, where the top row represents the highest rank.

Among the factors that influence the relative rank of various methods, the most important factor was heritability (*h*^2^) (Figure 3A). While all methods perform worse with increasing gene expression heritability, heritability disproportionately affects the performance of models that do not account for relatedness (i.e., negative binomial, Poisson and Linear), whereas when heritability is zero (*h*^2^ = 0), these approaches tend to perform slightly better. Therefore, for non-heritable genes, linear models perform slightly better than GEMMA, and negative binomial models work similarly or slightly better than MACAU. This observation most likely arises because linear and negative binomial models require fewer parameters and thus have a greater number of degrees of freedom. However, even in this setting, the difference between MACAU and negative binomial is small, suggesting that MACAU is robust to model misspecification and works reasonably well even for non-heritable genes. On the other hand, when heritability is moderate (*h*^2^ = 0.3) or high (*h*^2^ = 0.6), the methods that account for sample non-independence are much more powerful than the methods that do not. Because almost all genes are influenced by cis-eQTLs (47,48) and are thus likely heritable to some extent, MACAU’s robustness for non-heritable genes and its high performance gain for heritable genes make it appealing.

The second most important factor in relative model performance was the variation of total read counts across individuals (CV; Figure 3B). While all methods perform worse with increasing CV, CV particularly affects the performance of GEMMA. Specifically, when CV is small (0.3; as the baboon data), GEMMA works well and is the second best method behind MACAU. However, when CV is moderate (0.6) or high (0.9), the performance of GEMMA quickly decays: it becomes only the fourth best method when CV = 0.9. GEMMA performs poorly in high CV settings presumably because the linear mixed model fails to account for the mean-variance dependence observed in count data, which is in agreement with previous findings (60,104).

The other four factors we explored had small impacts on the relative performance of the alternative methods, although they did affect their absolute performance. For example, as one would expect, power increases with large effect sizes (PVE) (Figure S9) or large sample sizes (Figure S10), and decreases with large over-dispersion *σ*^2^ (Figure S11) or large *h_x_^2^* (Figure S12).

Finally, we included comparisons with edgeR (25) and DESeq2 (15). In the basic parameter simulation setting (*n* = 63, CV = 0.3, *h_x_^2^* = 0, PVE = 0.25, *h*^2^ = 0.3 and *σ*^2^ = 0.25), we again discretized the continuous predictor *x* into a binary 0/1 variable based on the median predictor value across individuals. Results for all methods are shown in Figure S13A. For the five methods also tested on a continuous predictor variable, the power on binarized values is much reduced compared with the power when the predictor variable is modeled without binarization (e.g. Figure S13A vs Figure 3). Further, neither edgeR nor DESeq2 perform well, consistent with the recent move from these methods towards linear models in differential expression analysis (3,7,46-48,105). This result is not contingent on having large sample sizes. In small sample size settings (n=6, n=10, and n=14, with samples balanced between the two classes, 0 or 1), MACAU again outperforms the other methods, though the power difference is much smaller (n=10 and n=14; Figures S13C and S31D) and sometimes negligible (n=6, Figure S13B).

In summary, the power of MACAU and other methods, as well as the power difference between methods, is influenced in a continuous fashion by multiple factors. Larger sample sizes, larger effect sizes, lower read depth variation, lower gene expression heritability, lower predictor variable heritability, and lower overdispersion all increase power. However, MACAU’s power is less diminished by high gene expression heritability and high read depth variability than the non-mixed model methods, while retaining the advantage of modeling the count data directly. In challenging data analysis settings (e.g., when sample size is low *and* effect size is low: Figure S13B for n=6), no method stands out, and using MACAU results in no or negligible gains in power relative to other methods. When the sample size is low (n=6) and effect sizes are large, however, MACAU consistently outperforms the other methods (n=6, Figure S14).

### Real Data Applications

To gain insight beyond simulation, we applied MACAU and the other six methods to three recently published RNAseq data sets.

The first data set we considered is the baboon RNAseq study (7) used to parameterize the simulations above. Expression measurements on 12,018 blood-expressed genes were collected by the Amboseli Baboon Research Project (ABRP) (71) for 63 adult baboons (26 females and 37 males), among which some were relatives. Here, we applied MACAU and the six other methods to identify genes with sex-biased expression patterns. Sex-associated genes are known to be enriched on sex chromosomes (106,107), and we use this enrichment as one of the criteria to compare method performance, as in (18). Because the same nominal *p*-value from different methods may correspond to different type I errors, we compared methods based on empirical false discovery rate (FDR). In particular, we permuted the data to construct an empirical null, estimated the FDR at any given *p*-value threshold, and counted the number of discoveries at a given FDR cutoff (see Methods and Materials for permutation details and a comparison between two different permutation procedures; Figure S15).

In agreement with our simulations, MACAU was the most powerful method of those we considered. Specifically, at an empirical FDR of 5%, MACAU identified 105 genes with sex-biased expression patterns, 40% more than that identified by the linear model, the second best method at this FDR cutoff (Figure 4A). At a more relaxed FDR of 10%, MACAU identified 234 sex-associated genes, 47% more than that identified by the negative binomial model, the second best method at this FDR cutoff (Figure 4A). Further, as expected, the sex-associated genes detected by MACAU are enriched on the X chromosome (the Y chromosome is not assembled in baboons and is thus ignored), and this enrichment is stronger for the genes identified by MACAU than by the other methods (Figure 4B). Of the remaining approaches, the negative binomial, linear model, and GEMMA all performed similarly and are ranked right after MACAU.

**Figure 4.**
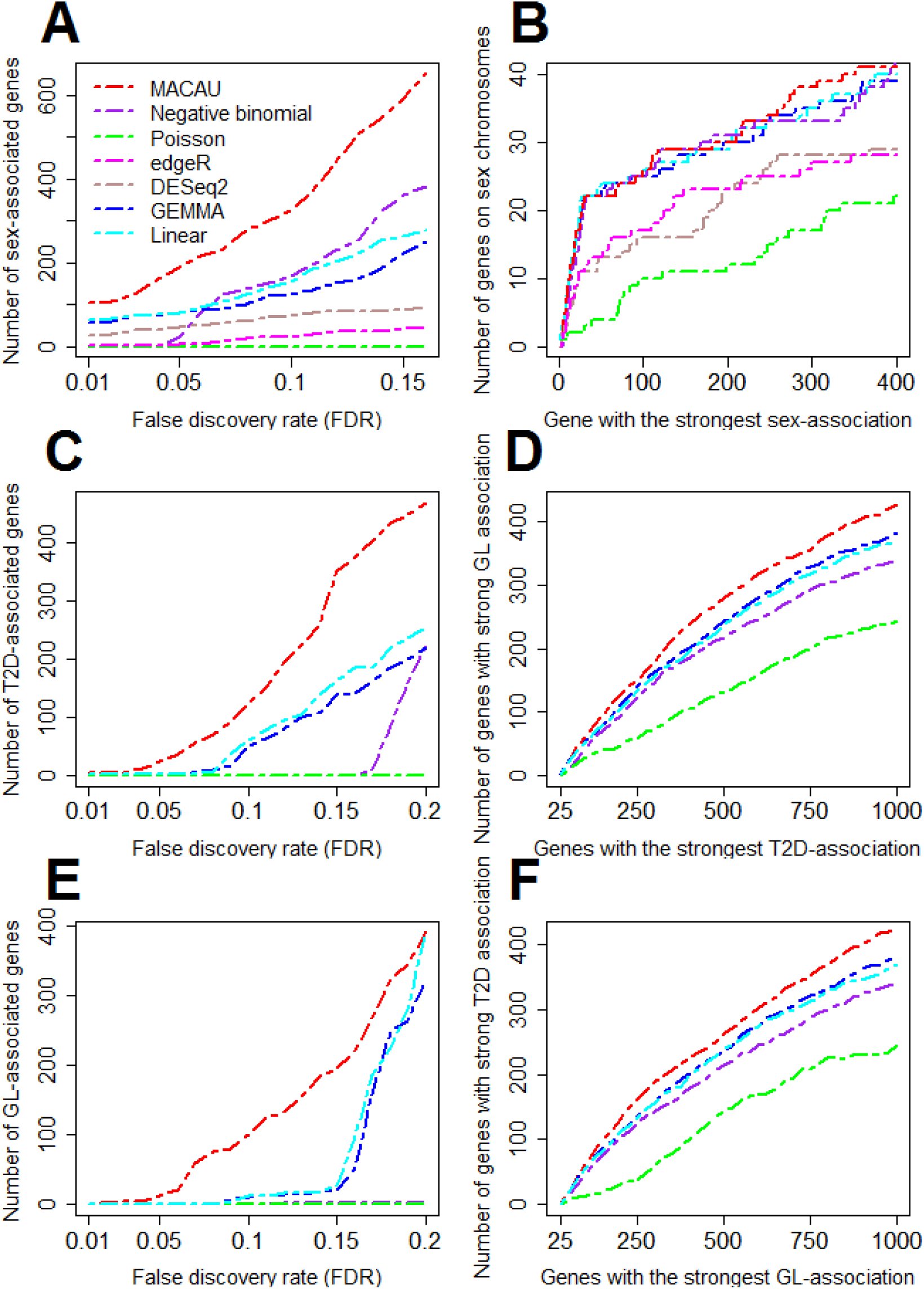
MACAU identifies more differentially expressed genes than other methods in the baboon (panels A and B) and FUSION (panels C, D, E, and F) data sets. Methods for comparison include MACAU (red), Negative binomial (purple), Poisson (green), edgeR (magenta), DESeq2 (rosybrown), GEMMA (blue), and Linear (cyan). (A) shows the number of sex-associated genes identified by different methods at a range of empirical false discovery rates (FDRs). (B) shows the number of genes that are on the X chromosome out of the genes that have the strongest sex association for each method (note that the Y chromosome is not assembled in baboons and is thus ignored). For instance, in the top 400 genes identified by MACAU, 41 of them are also on the X chromosome. (C) shows the number of T2D-associated genes identified by different methods at a range of empirical false discovery rates (FDRs). (D) shows the number of genes that are in the list of top 1,000 genes most significantly associated with GL out of the genes that have the strongest association for T2D for each method. For instance, in the top 1,000 genes with the strongest T2D association identified by MACAU, 428 of them are also in the list of top 1,000 genes with the strongest GL association identified by the same method. (E) shows the number of GL-associated genes identified by different methods at a range of FDRs. (F) shows the number of genes that are in the list of top 1,000 genes most significantly associated with T2D out of the genes that have the strongest association for GL for each method. T2D: type II diabetes; GL: fasting glucose level.

**Table 1.** Current approaches for identifying differentially expressed genes in RNAseq.

| Statistical method | Directly models counts? | Controls for biological covariates? | Controls for sample non-independence? | Example software that implements the method |
| --- | --- | --- | --- | --- |
| Linear regression | No | Yes | No | R and many others |
| Linear mixed model | No | Yes | Yes | GEMMA (9) and EMMA (134) |
| Poisson model | Yes | Some methods do | No | GLMP (135) and DEGseq (20) |
| Negative binomial model | Yes | Some methods do | No | edgeR (25), DESeq (15) and GLMNb(135) |
| Poisson mixed model | Yes | Yes | Yes | MACAU |

The Poisson model performs the worst, and edgeR and DESeq2 fall between the Poisson model and the other methods (Figures 4A and 4B).

The second data set we considered is an RNAseq study on type II diabetes (T2D) collected as part of the Finland-United States Investigation of NIDDM Genetics (FUSION) Study (61). Here, the data were collected from skeletal muscle samples from 267 individuals with expression measurements on 21,753 genes. Individuals are from three municipalities (Helsinki, Savitaipale, and Kuopio) in Finland. Individuals within each municipality are more closely related than individuals between municipalities (e.g., the top genotype principal components generally correspond to the three municipalities; Figure S16). Two related phenotypes were available to us: 162 individuals with T2D or NGT (normal glucose tolerance) status (i.e., case/control) based on the oral glucose tolerance test (OGTT) and 267 individuals with the quantitative trait fasting glucose level (GL), a biologically relevant trait of T2D.

We performed analyses to identify genes associated with T2D status as well as genes associated with GL. To accommodate edgeR and DESeq2, we also discretized the continuous GL values into binary 0/1 categories based on the median GL value across individuals. We refer to the resulting values as GL01. Therefore, we performed two sets of analyses for GL: one on the continuous GL values and the other on the discretized GL01 values. Consistent with simulations and the baboon data analysis, MACAU identified more T2D-associated genes and GL-associated genes than other methods across a range of empirical FDR values. For the T2D analysis, MACAU identified 23 T2D-associated genes at an FDR of 5%, while GEMMA and the linear model, the second best methods at this FDR cutoff, identified only 1 T2D-associated gene (Figure 4C). Similarly, at an FDR of 10%, MACAU identified 123 T2D-associated genes, 51% more than that identified by the linear model, the second best method at this FDR cutoff (Figure 4C). For GL analysis, based on an FDR of 5%, MACAU detected 12 DE genes, while the other methods did not identify any DE genes at this FDR cutoff. At an FDR of 10%, MACAU identified 100 GL associated genes, while the second best methods -- the linear model and GEMMA -- identified 12 DE genes (Figure 4E). For the dichotomized GL01, none of the methods detected any DE genes even at a relaxed FDR cutoff of 20%, highlighting the importance of modeling the original continuous predictor variable in DE analysis.

Several lines of evidence support the biological validity of the genes detected by MACAU. First, we performed Gene Ontology (GO) analysis using LRpath (108) on T2D and GL associated genes identified by MACAU, as in the FUSION study (61) (Figure S17). The GO analysis results for T2D and GL are consistent with previous studies (61,109) and are also similar to each other, as expected given the biological relationship between the two traits. In particular, T2D status and high GL are associated with decreased expression of cellular respiratory pathway genes, consistent with previous observations (61,109). T2D status and GL are also associated with several pathways that are related to mTOR, including generation of precursor metabolites, poly-ubiquitination and vesicle trafficking, in agreement with a prominent role of mTOR pathway in T2D etiology (110-113).

Second, we performed overlap analyses between T2D and GL associated genes. We reasoned that T2D-associated genes are likely associated with GL because T2D shares a common genetic basis with GL (114-116) and T2D status is determined in part by fasting glucose levels. Therefore, we used the overlap between genes associated with T2D and genes associated with GL as a measure of method performance. In the overlap analysis, genes with the strongest T2D association identified by MACAU show a larger overlap with the top 1,000 genes that have the strongest GL association than did genes identified by other methods (Figure 4D). For instance, among the top 100 genes with the strongest T2D-association evidence from MACAU, 63 of them also show strong association evidence with GL. In contrast, only 55 of the top 100 genes with the strongest T2D-association identified by GEMMA, the second best method, show strong association evidence with GL. We observed similar results, with MACAU performing the best, when performing the reciprocal analysis (overlap between genes with the strongest GL-association and the top 1,000 genes that have the strongest T2D-association: Figure 4F). To include the comparison with edgeR and DESeq2, we further examined the overlap between T2D associated genes and GL01 associated genes for all methods (Figure S18). Again, MACAU performs the best, followed by GEMMA and the linear model, and neither edgeR nor DESeq2 perform well in this context (Figure S18). Therefore, MACAU appears to both confer more power to identify biologically relevant DE genes and be more consistent across analyses of related phenotypes.

To assess the type I error rate of various methods, we permuted the trait data from the baboon and the FUSION studies. Consistent with our simulation results, the *p*-values from MACAU and GEMMA under the permuted null were close to uniformly distributed (slightly conservative) in both data sets, whereas the other methods were not (Figures S19 and S20). In addition, none of the methods compared here are sensitive to outliers in the two data sets (Figures S21-S23).

Finally, although large, population-based RNAseq data sets are becoming more common, MACAU’s flexible PMM modeling framework allows it to be applied to DE analysis in small data sets with unrelated individuals as well. In this setting, MACAU can use the gene expression covariance matrix as the ***K*** matrix to control for hidden confounding effects that are commonly observed in sequencing studies (49-52). Hidden confounders can induce similarity in gene expression levels across many genes even though individuals are unrelated (53-57), similar to the effects of kinship or population structure. Therefore, by defining ***K*** using a gene expression (instead of genetic) covariance matrix, MACAU can effectively control for sample non-independence induced by hidden confounders, thus extending the linear mixed model widely used to control for hidden confounders in array based studies (53-57) to sequencing count data.

To illustrate this application, we analyzed a third data set on lymphoblastoid cell lines (LCLs) derived from 69 unrelated Nigerian individuals (YRI) (3) from the HapMap project (117), with expression measurements on 13,319 genes. We also aimed to identify sex-associated genes in this data set. To demonstrate the effectiveness of MACAU in small samples, we randomly subsampled individuals from the data to create small data sets with either *n* = 6 (3 males and 3 females), *n* = 10 (5 males and 5 females), or *n* = 14 individuals (7 males and 7 females). For each sample size *n*, we performed 20 replicates of random subsampling and then evaluated method performance by averaging across replicates. In each replicate, we used the gene expression covariance matrix as ***K*** and compared MACAU’s performance against other methods. Because of the small sample size, none of the methods were able to identify DE genes at an FDR cutoff of 10%, consistent with recent arguments that at least 6-12 biological replicates are needed to ensure sufficient power and replicability in DE analysis (11). We therefore used enrichment of genes on the sex chromosomes to compare the performance of different methods (Figure S24). The enrichment of top ranked sex-associated genes on sex chromosomes has previously been used for method comparison and is especially suitable for comparing methods in the presence of batch effects and other hidden confounding factors (118).

In this comparison, MACAU performs the best of all methods when the sample size is either *n* = 10 or *n* = 14, and is ranked among the best (together with the negative binomial model) when *n* = 6. For instance, when *n* = 6, among the top 50 genes identified by each method, the number of genes on the sex chromosomes for MACAU, negative binomial, Poisson, edgeR, DESeq2, GEMMA, and Linear are 3.3, 2.7, 3.1, 1.8, 3.0, 2.0, and 2.4, respectively. The advantage of MACAU becomes larger when the sample size increases: for example, when *n* = 14, an average of 10.6 genes in the top 50 genes from MACAU are on the sex chromosomes, which is again larger than that from the negative binomial (8.3), Poisson (6.0), edgeR (6.65), DESeq2 (8.8), GEMMA (9.8), or Linear (8.05). These results suggest that MACAU can also perform better than existing methods in relatively small sample study designs with unrelated individuals by controlling for hidden confounders. However, MACAU’s power gain is much smaller in this setting than in the first two data sets we considered (the baboon and Fusion data). In addition, MACAU’s power gain is negligible in the case of n=6 when compared with the second best method, though its power gain over the commonly used edgeR and DESeq2 is still substantial. MACAU’s small power gain in this data presumably stems from both the small sample size and the small effect size of sex in the data, consistent with previous reports for blood cell-derived gene expression (3,7,119).

## Discussion

Here, we present an effective Poisson mixed effects model, together with a computationally efficient inference method and software implementation in MACAU, for identifying DE genes in RNAseq studies. MACAU directly models count data and, using two random effects terms, controls for both independent over-dispersion and sample non-independence. Because of its flexible modeling framework, MACAU controls for type I error in the presence of individual relatedness, population structure, and hidden confounders, and MACAU achieves higher power than several other methods for DE analysis across a range of settings. In addition, MACAU can easily accommodate continuous predictor variables and biological or technical covariates. We have demonstrated the benefits of MACAU using both simulations and applications to three recently published RNAseq data sets.

MACAU is particularly well-suited to data sets that contain related individuals or population structure. Several major population genomic resources contain structure of these kinds. For example, the HapMap population (117), the Human Genome Diversity Panel (120), the 1000 Genomes Project in humans (121) as well as the 1001 Genomes Project in Arabidopsis (122) all contain data from multiple populations or related individuals. Several recent large-scale RNAseq projects also collected individuals from genetically differentiated populations (46). MACAU is also well-suited to analyzing genes with moderate to high heritability. Previous studies in humans have shown that, while heritability varies across genes, many genes are moderately or highly heritable, and almost all genes have detectable eQTL (47,123). Analyzing these data with MACAU can reduce false positives and increase power. Notably, even when genes exhibit zero heritability, our results show that MACAU incurs minimal loss of power compared with other approaches.

While we have mainly focused on illustrating the benefits of MACAU for controlling for individual relatedness and population stratification, we note that MACAU can be used to control for sample non-independence occurred in other settings. For example, cell type heterogeneity (55) or other hidden confounding factors (53) are commonly observed in sequencing studies and can induce gene expression similarity even when individuals are unrelated (49-52). Because the gene expression covariance matrix ***K*** contains information on sample non-independence caused by hidden confounding factors (53-57), MACAU could be applied to control for hidden confounding effects by using the gene expression covariance as the ***K*** matrix. Therefore, MACAU provides a natural avenue for extending the commonly used mixed effects model for controlling for hidden confounding effects (53-56) in array-based studies to sequencing studies. In addition, although we have designed MACAU for differential expression analysis, we note that MACAU may also be effective in other common settings. For example, MACAU could be readily applied in QTL mapping studies to identify genetic variants that are associated with gene expression levels estimated using RNAseq or related high-throughput sequencing methods.

In the present study, we have focused on demonstrating the performance of MACAU in three published RNAseq data sets with sample sizes ranging from small (n=6) to medium (n=63) to large (n=267), relative to the size of most current RNAseq studies. Compared with small sample studies, RNAseq studies with medium or large sample sizes are better powered and more reproducible, and are thus becoming increasingly common in genomics (10,11). For example, a recent comparative study makes explicit calls for medium to large sample RNAseq studies performed with at least 12 replicates per condition (i.e. n>=24) (11). However, we recognize that many RNAseq studies are still carried out with a small number of samples (e.g. 3 replicates per condition). As our simulations make clear, the power of all analysis methods is dramatically reduced with decreasing sample size, conditional on fixed values of other factors that influence power (e.g., effect size, gene expression heritability). Thus, MACAU’s advantage is no longer obvious in simulated data with only 3 replicates per condition when the effect size is also small (although its advantage becomes apparent when the simulated effect size increases: Figures S13B and S14). In addition, MACAU’s advantage is much smaller and sometimes negligible in the small real data set when compared with the medium and large data sets analyzed here. Furthermore, because MACAU requires estimating one more parameter than other existing methods, MACAU requires at least five samples to run while existing DE methods require at least four. Therefore, MACAU may not confer benefits to power in some settings, and is especially likely (like all methods) to be underpowered in very small sample sizes with small effect sizes. Future extensions of MACAU are likely needed to ensure its robust performance in small as well as moderate to large samples. For example, further power improvements could be achieved by borrowing information across genes to estimate the over-dispersion parameter (15,22,25) or building in a hierarchical structure to model many genes at once.

Like other DE methods (24,25), MACAU requires data pre-processing to obtain gene expression measurements from raw sequencing read files. This data pre-processing step may include read alignment, transcript assembly, alternative transcript quantification, transcript measurement, and normalization. Many methods are available to perform these tasks (12,14,68,124-129) and different methods can be differentially advantageous across settings (68,124,130). Importantly, MACAU can be paired with any pre-processing method that retains the count nature of the data. While we provide a preliminary comparison of several methods here (see Materials and Methods; Figure S3), a full analysis of how different data pre-processing choices affect MACAU’s performance in alternative study designs is beyond the scope of this paper. Notably, recent results suggest that a recommended approach is to incorporate data pre-processing and DE analysis into the same, joint statistical framework (131), which represents an important next step for the MACAU software package.

We note that, like many other DE methods (15,25), we did not model gene length in MACAU. Because gene length does not change from sample to sample, it does not affect differential expression analysis on any particular gene (15,25). However, gene length will affect the power of DE analysis across different genes: genes with longer length tend to have a larger number of mapped reads and more accurate expression measurements, and as a consequence, DE analysis on these genes tends to have higher statistical power (2,70,132). Gene length may also introduce sample-specific effects in certain data sets (133). Therefore, understanding the impact of, and taking into account gene length effects, in MACAU DE analysis represents another possible future extension.

Currently, despite the newly developed computationally efficient algorithm, applications of MACAU can still be limited by its relatively heavy computational cost. The MCMC algorithm in MACAU scales quadratically with the number of individuals/samples and linearly with the number of genes. Although MACAU is two orders of magnitude faster than the standard software MCMCglmm for fitting Poisson mixed effects models (Table S1), it can still take close to 20 hours to analyze a data set of the size of the FUSION data we considered here (267 individuals and 21,753 genes). Therefore, new algorithms will be needed to use MACAU for data sets that are orders of magnitude larger.

## URLs

The software implementation of MACAU is freely available at: www.xzlab.org/software.html.

## Competing Interests

The authors declare that they have no competing interests.

## Funding

This study was supported by NIH fund R01HG009124. XZ is also supported by R01HL117626 (PI Abecasis), R21ES024834 (PI Pierce), R01HL133221 (PI Smith), and a grant from the Foundation for the National Institutes of Health through the Accelerating Medicines Partnership (BOEH15AMP, co-PIs Boehnke and Abecasis). JT is supported by 1R01GM102562 and R21AG049936. LS is supported by U01DK062370 (PI Boehnke). SS is supported by a scholarship from the China Scholarship Council.

## Acknowledgements

We thank Matthew Stephens for insight and support on previous versions of MACAU. We thank Baylor College of Medicine Human Genome Sequencing Center for access to the current version of the baboon genome assembly (Panu 2.0). We thank FUSION investigators for access to the FUSION expression data.

